# Parallel rapid phenotypic differentiation along climatic gradients in multiple introduced lineages of an alien plant species

**DOI:** 10.1101/2024.05.13.593839

**Authors:** Saeko Matsuhashi, Hiroshi Kudoh, Michio Oguro, Satoki Sakai

**Author notes:** Author for correspondence: Saeko Matsuhashi.

## Abstract

Widespread plant species are faced with various climatic gradients and undergo clinal differentiation. Alien species which have been introduced repeatedly can be used to investigate the parallel processes of clinal differentiation during their range expansions. *Cardamine hirsuta* L. has been unintentionally introduced to Japan at least three times and has become widely distributed across Japan. To elucidate the processes of clinal differentiation, we conducted a common garden experiment using three lineages of *C. hirsuta* (North, East, and West). We tested for phenotypic differentiation among and within the lineages, and for a common pattern of clinal variations in phenotypic traits. We detected differentiations of flowering phenology and body size within and among the lineages. The East lineage showed delayed flowering phenology and larger mass, whereas the West lineage showed rapid flowering phenology and smaller mass. In addition, three patterns of variation between climate factors in seed source habitats and phenotypic traits were detected in all three lineages: higher temperature was related to earlier flowering and lower leaf number at the start of flowering, and more hours of sunshine were related to shorter flowering period. These patterns imply independent and parallel differentiations in the three lineages. These results suggest that analyses of distinct lineages allow repeated observations of differentiation processes and rapid responses of species during successful distribution in new environments.

## Introduction

Widespread plant species are faced with various climatic gradients such as temperature, rainfall, snowfall, and sunlight conditions, and must adapt to local environments (Joshi et al. 2001; Petrů et al. 2006). Understanding which climatic factors act as selective pressures and how clinal differentiation is constructed allows us to estimate how plant distributions are determined and to understand evolution during range expansions in the past, present, and future (Li et al. 1998; Olsson and Agren 2002; Paccard et al. 2014; Preite et al. 2015). Alien species often provide opportunities to investigate clinal differentiation during their range expansions (Hulme 2011; Colautti and Barrett 2013; Liao et al. 2016; Helsen et al. 2020) because adaptation to novel climatic conditions in introduced areas is essential for their success. Clinal differentiation in adaptive traits along climatic gradients has been reported in several widespread alien species (Weber and Schmid 1998; Kollmann and Bañuelos 2004; Maron et al. 2004; Novy et al. 2013; Oduor et al. 2016). For example, flowering initiation time and size at flowering have differentiated along latitude in introduced *Lythrum salicaria* (Montague et al. 2008), and height and aboveground biomass have differentiated along altitudinal gradients in introduced *Senecio inaequidens* (Monty and Mahy 2009).

However, structures of genetic variations governing phenotypic traits in alien species should be interpreted carefully because they may be affected by multiple introductions (van Boheemen et al. 2019). Introductions of distinct lineages of the same alien species into non-native ranges from multiple different sources have been reported (Bossdorf et al. 2005; Dlugosch and Parker 2008), and lineages with different origins may have different sets of genetic variations affecting the same traits. By taking lineage composition into account, we can distinguish whether clinal differentiation resulted from differentiation within lineages or from different distributions of multiple lineages. However, few studies of alien species have examined clinal differentiation of phenotypic traits and distributions of lineages simultaneously.

*Cardamine hirsuta* L. (Brassicaceae), a winter annual native to Europe, has been introduced to Japan at least three times and has become widely distributed across Japan (Matsuhashi et al. 2016). The three lineages can be distinguished clearly in the current Japanese populations using genetic markers, suggesting that intercrossing between different lineages has been infrequent (Matsuhashi et al. 2016). This situation allowed us to study clinal differentiation within and between the lineages. Moreover, we could test whether there is a common pattern of genetic differentiation affecting phenotypic traits along climatic gradients during the range expansions of the three lineages.

Here, we examined whether genetic differentiations in growth-related and phenology traits along climatic gradients have shaped Japanese populations of *C. hirsuta*. For this purpose, we conducted a common garden experiment using materials collected from a wide range of geographic regions across Japan. We analyzed how genetic variation of the traits correlated with climatic factors for the distinct lineages previously identified using neutral genetic markers (Matsuhashi et al. 2016). We asked the following questions: (i) Are there genetic differentiations in growth-related and phenological traits among the three lineages? (ii) Are there clinal differentiations of traits within each lineage? (iii) Are there shared patterns in the clinal differentiations of traits among the three lineages? By answering these questions, we identified traits that have genetically differentiated in response to climatic gradients and the climatic factors that are expected to have driven clinal genetic differentiation during range expansions of introduced *C. hirsuta*.

## Materials and Methods

### Plant species

*C. hirsuta* is native to Europe, but is distributed worldwide today; it is found in North America, Australia, North Africa, Asia, and other areas (Lihova et al. 2006). In Japan, the earliest specimen was collected in 1974 (Kudoh et al. 1992), and its distribution has expanded recently (Kudoh et al. 2007). Three distinct lineages were detected using microsatellite analyses (Matsuhashi et al. 2016). They are widely distributed in northern, eastern, and western Japan, and are referred to as the North, East, and West lineages, respectively. The species shows seed dormancy during summer and germination begins in September (Kudoh et al. 2007). The flowers are self-compatible and the flowering period extends from February to April (Yatsu et al. 2003; Kudoh et al. 2007).

### Sample collection

We collected seeds for growth experiments from 92 plants (maternal families) from 57 sites in Japan (Fig. 1). When multiple samples were collected from a single site, seeds were collected from plants growing at least 5 m apart. The samplings were conducted from March to May in 2009 and 2010. All sampling sites were located within ten kilometers of the nearest meteorological stations operated by the Japan Meteorological Agency. From all the corresponding stations, we obtained meteorological data on temperature, precipitation, and hours of sunlight for 1999–2009. To estimate the climate conditions of each site during the growth period of *C. hirsuta* (from September to the next April; Kudoh et al. 2007), we calculated the mean daily temperature, the total precipitation, and the total hours of sunshine from September to the next April.

**Fig. 1:**
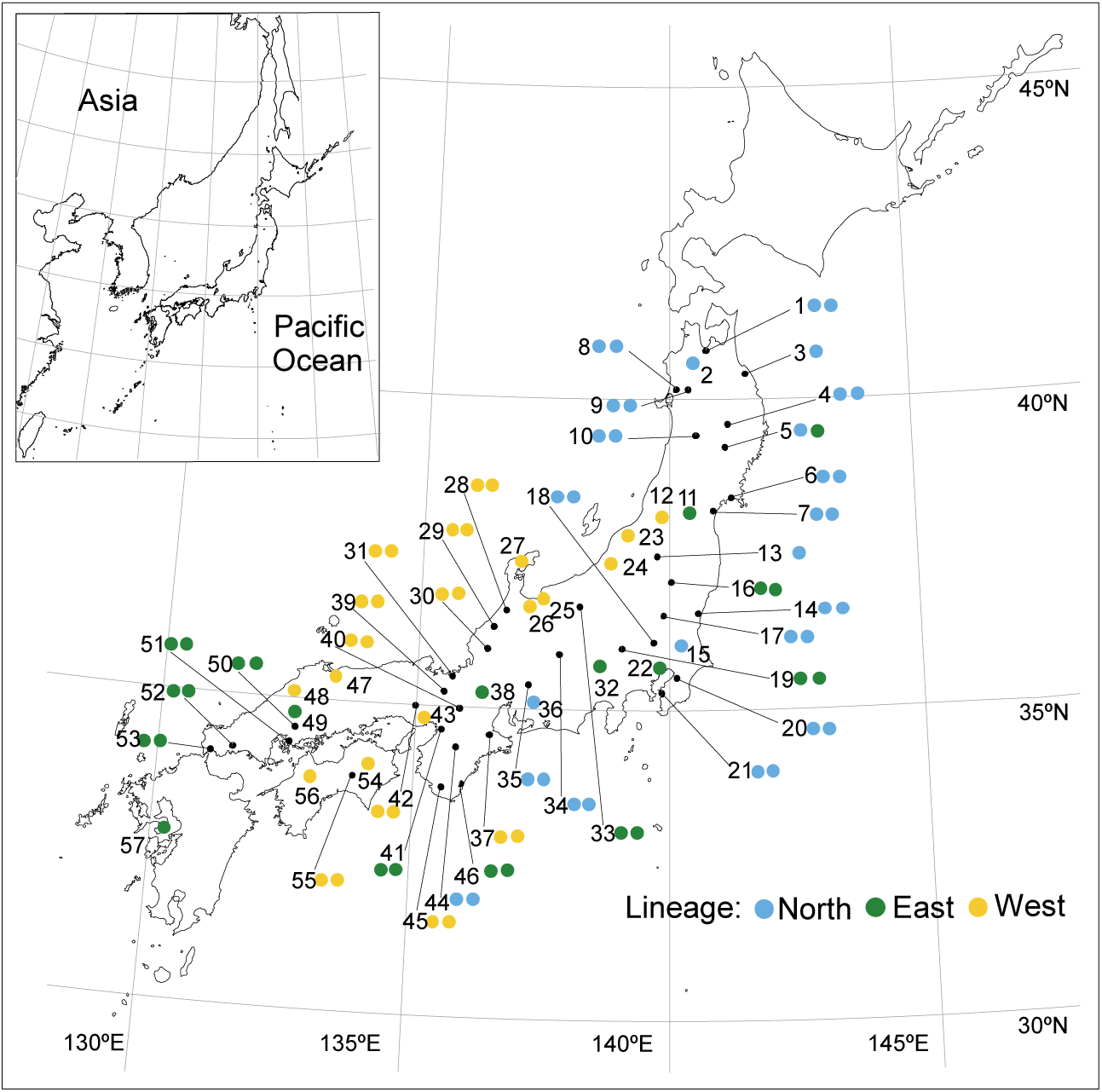
The locations of 57 sample collection sites where maternal plants were collected. The color of each circle symbol indicates the lineage to which the maternal plant belongs. The number of circle symbols indicates the number of maternal plants collected at a site.

All maternal plants had been genotyped using nine microsatellite markers as described by Matsuhashi *et al*. (2016). According to the results of Bayesian clustering, 35, 25 and 32 maternal families were assigned to the North, East, and West lineages, respectively (Fig. 1). As the measures of genetic differences among the maternal families, we used the values calculated in our previous study (Matsuhashi et al. 2016), i.e., PC1 and PC2 from the result of a principal coordinate analysis (PCoA) on D*_A_* genetic distance (Nei et al. 1983).

### Phenology monitoring and trait measurements

To compare reproductive, vegetative, and phenological traits among families, we conducted a growth experiment from October 2011 to March 2012 in a controlled environment. For each maternal family, 20 seeds were sown on perlite in a petri dish. The dishes were placed in an incubator under 20°C/10°C of day/night temperature with 12 h day length and kept moist with distilled water throughout these periods. The seeds germinated 1-3 weeks after sowing. Four weeks after sowing, each seedling was transplanted to a plastic pot (7.5cm diameter) containing a soil mixture of vermiculite:perlite:culture soil = 3:1:1 (0.4gN, 1.0gP, and 0.6gK per kg culture soil) by volume. Four weeks after transplantation, we selected 3 plants with 5-8 true leaves from each maternal family and exposed them to low temperature (5°C) for 3 weeks under darkness for vernalization. They were then transferred to a phytotron (NCP-1.5, NK system, Japan) and grown under 20°C/10°C of day/night temperature with 12 h day length.

For each plant, the number of leaves in the rosette was counted at the end of the vernalization treatment. Then, the growth of 276 plants (2-3 plants per maternal family after removing several plants which were accidentally damaged) was monitored every day after transfer to the growth chamber. We recorded the dates of opening of the first and last flowers of the main inflorescence of each plant. We calculated the number of days from the end of the vernalization treatment to the opening of the first flower (days to flowering after vernalization), and from the opening of the first flower to the opening of the last flower of the main inflorescence (flowering period).

After all flowers of the main inflorescence had finished flowering, we recorded the number of flowers, including those on the lateral inflorescences, and the number of rosette leaves, including withered ones, for each plant. We randomly selected four fruits on the main inflorescence and counted the number of ovules to compare potential seed production ability. Roots, rosette leaves, and reproductive parts including supporting organs (inflorescences with stem and cauline leaves) of each plant were separated and dried in an oven at 60°C for 3 days. Root dry mass, rosette leaf dry mass, and reproductive dry mass were measured and recorded. Total plant dry mass was also calculated by adding the dry masses of all parts.

We thus obtained the following ten phenotypic traits for each plant: a) days to flowering after vernalization, b) flowering period, c) flower number, d) leaf number when the first flower opened, e) leaf number when the last flower opened, f) root dry mass, g) above-ground dry mass, h) reproductive dry mass, i) total plant dry mass, and j) total number of ovules in four fruits.

### Analyses of the effects of lineage, genetic distance, and climate on the plant traits

All the analyses described below were conducted using R 3.2.5 (R Core Team 2016) unless otherwise noted.

To elucidate the genetic differentiation of the phenotypic traits among the lineages and among the maternal families, we conducted nested analysis of variance (nested-ANOVA) in which the maternal families were nested within the three lineages.

To evaluate the dependencies of the phenotypic values on the lineages, the climatic conditions of the sampling sites, and the genetic distances, generalized linear mixed models (GLMMs) were applied. The response variables of the models were the ten phenotypic traits. The lineage, the climatic conditions of the sampling sites (mean daily temperature, total precipitation, and total hours of sunshine from September to April), and the measures of genetic distances (PC1 and PC2 of the results of PCoA) were used as the explanatory variables. To deal with non-linear relationships, quadratic terms of the climatic conditions and the measures of genetic distances were also included in the models. To detect differences in the relationships of the phenotypic traits to the climatic conditions and to the measures of genetic distance among lineages, interaction terms between the lineage and the climatic conditions and between the lineage and the measures of genetic distances were included. The identities of maternal families nested within the identities of sampling sites were used as the random effect.

Implementation of the GLMMs was conducted using the *MCMCglmm* function in the *MCMCglmm* package (Hadfield 2010). For the error distributions of the models, normal distributions were used for the continuous traits including the days to flowering after vernalization and the flowering period (they are continuous by nature) and Poisson distributions were used for the discrete traits. For the prior distributions of model parameters, uninformative priors were used: the default parameters were used for the fixed effect (B-structure), an inverse-Gamma distribution (*V* = 1 and *nu* = 0.002) was used for the variance (R-structure), and a scaled Fisher F distribution (*V* = 1, *nu* = 1, *alpha.mu* = 0, and *alpha.V* = 10^9^) was used for the random effects (G-structure) (de Villemereuil et al. 2013). A Markov chain Monte Carlo (MCMC) sampler with 130,000 iterations, 30,000 burn-in, and 100 thinning interval was used to estimate the posterior probability distribution of the model parameters. Convergences of the models were checked by Gelman and Rubin’s convergence diagnostic (Gelman and Rubin 1992) using the *gelman.diag* function in the *coda* package (Plummer et al. 2006). If a MCMC sampler did not converge, values of iterations, burn-in, and thinning interval were doubled, and the analysis was run again. This procedure was repeated until the MCMC converged. Before the analyses, values of climatic conditions and measures of genetic distance were scaled to mean = 0 and standard deviation = 1 using the *scale* function in R to improve mixing of the MCMCs and to obtain standardized regression coefficients.

To obtain the models having the best predictive ability, combinations of the explanatory variables were selected to minimize deviance information criteria (DIC) (Spiegelhalter et al. 2002) by stepwise model selection procedures. We tried both backward and forward model selection procedures and selected the model having the smaller value of DIC as the model with the best predictive ability (best model).

## Results

### Phenotypic differences among the lineages

In the experiment using the 92 maternal families, we found differences among the three lineages for most traits (Table 1, Fig. 2, 3). The results of nested-ANOVA (Table 1) showed that all the measured traits except for the total number of ovules differed significantly among the lineages and among the maternal families within the lineages. For all traits, the effect sizes *η*^2^ of the maternal families within the lineages were larger than those of the lineages (Table 1). The East lineage showed delayed flowering (mean days to flowering after vernalization: 45.3 days) and longer flowering period (mean: 21.5 days), while the West lineage tended to show earlier flowering (mean: 36.5 days) and shorter flowering period (mean: 17.1 days) than the other two lineages (Fig. 2a, b). Total mass and root mass were larger in the East lineage than in the other two lineages (Fig. 2i, f). Total above-ground, root,and reproductive mass were smaller in the West lineage than in the other two lineages (Fig. 2g, h, i). In contrast, the flower number was greater in the West lineage than in the other two lineages (Fig. 2c). The East lineage had larger ranges in the leaf number when the first flower of each plant opened and the leaf number when the last flower opened than the other lineages (Fig. 2d, e). The total number of ovules was significantly different among the maternal families within all the lineages, but not among the lineages (Table 1).

**Fig. 2:**
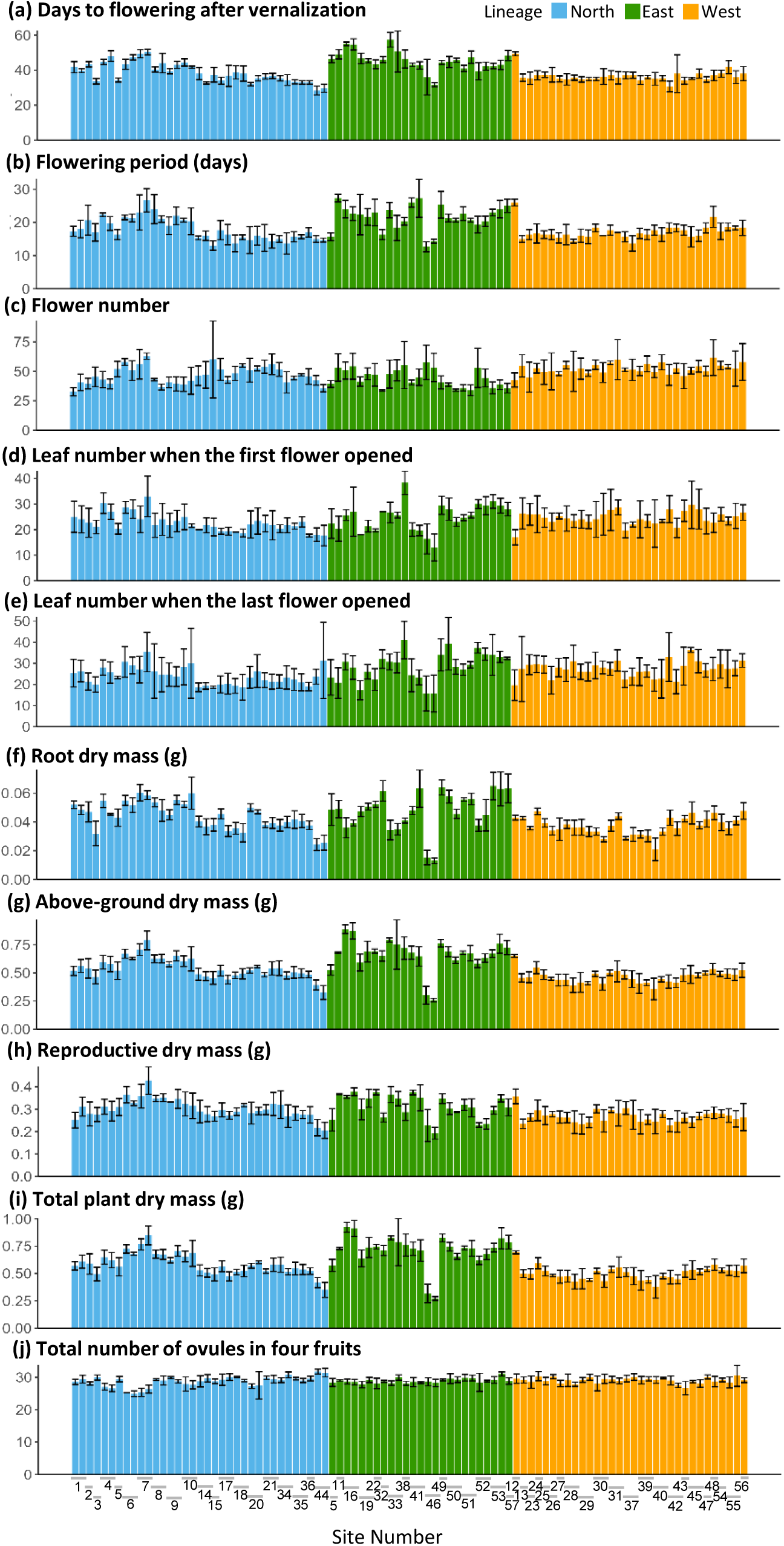
The measured values of ten traits (a∼j). Each bar indicates the mean of each maternal family. The bars are sorted by the three lineages (North, East, and West) and arranged in order of the site numbers in Fig. 1.

**Table 1:** The results of nested-ANOVA. The differentiation of the ten phenotypic traits among the lineages (North, East, and West) and among the maternal families within each lineage. The effect size *η^2^* is calculated from the sum of squares of the effect divided by the total sum of squares.

| Traits | Source of Variation | Sum of Squares | Degrees of Freedoms | Mean Square | $\eta^2$ | $F$ | $P$ |
| --- | --- | --- | --- | --- | --- | --- | --- |
| Days to flowering after vernalization | Lineage | 3476 | 2 | 1737.8 | 0.296 | 164.49 | <0.001 |
|  | Lineage*Maternal family | 6313 | 89 | 70.9 | 0.538 | 6.71 | <0.001 |
|  | Residuals | 1944 | 184 | 10.6 |  |  |  |
| Flowering period | Lineage | 844.4 | 2 | 422.2 | 0.190 | 73.72 | <0.001 |
|  | Lineage*Maternal family | 2604.7 | 89 | 29.3 | 0.585 | 5.11 | <0.001 |
|  | Residuals | 1002.3 | 175 | 5.7 |  |  |  |
| Flower number | Lineage | 2777 | 2 | 1388.3 | 0.100 | 17.87 | <0.001 |
|  | Lineage*Maternal family | 11289 | 89 | 126.8 | 0.405 | 1.63 | <0.01 |
|  | Residuals | 13832 | 178 | 77.7 |  |  |  |
| Leaf number when the first flower opened | Lineage | 266 | 2 | 133.15 | 0.034 | 7.08 | <0.01 |
|  | Lineage*Maternal family | 4075 | 89 | 45.79 | 0.528 | 2.44 | <0.001 |
|  | Residuals | 3384 | 180 | 18.8 |  |  |  |
| Leaf number when the last flower opened | Lineage | 997 | 2 | 498.3 | 0.066 | 12.10 | <0.001 |
|  | Lineage*Maternal family | 6428 | 89 | 72.2 | 0.428 | 1.75 | <0.001 |
|  | Residuals | 7581 | 184 | 41.2 |  |  |  |
| Root dry mass | Lineage | 0.005 | 2 | 0.002 | 0.131 | 85.85 | <0.001 |
|  | Lineage*Maternal family | 0.026 | 89 | 0.000 | 0.729 | 10.71 | <0.001 |
|  | Residuals | 0.005 | 182 | 0.000 |  |  |  |
| Above-ground dry mass | Lineage | 1.563 | 2 | 0.781 | 0.317 | 154.61 | <0.001 |
|  | Lineage*Maternal family | 2.450 | 89 | 0.028 | 0.497 | 5.45 | <0.001 |
|  | Residuals | 0.920 | 182 | 0.005 |  |  |  |
| Reproductive dry mass | Lineage | 0.085 | 2 | 0.042 | 0.109 | 31.67 | <0.001 |
|  | Lineage*Maternal family | 0.452 | 89 | 0.005 | 0.579 | 3.80 | <0.001 |
|  | Residuals | 0.243 | 182 | 0.001 |  |  |  |
| Total plant dry mass | Lineage | 1.734 | 2 | 0.867 | 0.327 | 235.27 | <0.001 |
|  | Lineage*Maternal family | 2.903 | 89 | 0.033 | 0.547 | 8.85 | <0.001 |
|  | Residuals | 0.667 | 181 | 0.004 |  |  |  |
| Total number of ovules in four fruits | Lineage | 74 | 2 | 36.85 | 0.007 | 1.51 | 0.224 |
|  | Lineage*Maternal family | 5805 | 89 | 65.22 | 0.570 | 2.67 | <0.001 |
|  | Residuals | 4302 | 176 | 24.44 |  |  |  |

### Effects of climate factors and genetic differences within each lineage on the traits

In the GLMM results, the lineages, the genetic distance (PC1 and/or PC2) and at least one of the climate factors were selected as the fixed effects for all traits. The relationships between a trait and a climate factor, and between a trait and genetic distance were different among the lineages for most of the 50 relationships ((3 climate factors + 2 genetic axes) x 10 traits) analyzed. For example, with increasing hours of sunshine, flower number increased in the North lineage, decreased in the East lineage, and showed no change in the West lineage. However, the three relationships between plant traits and climate factors were common to all lineages: days to flowering after vernalization (Fig. 3a, *p* < 0.001) and leaf number when the first flower opened (Fig. 3b, *p* = 0.006) decreased with increasing temperature, and flowering period (Fig. 3c, *p* = 0.012) decreased with increasing hours of sunshine. A positive relationship between root dry mass (*p* = 0.004) and PC2 was also detected in all lineages.

**Fig. 3:**
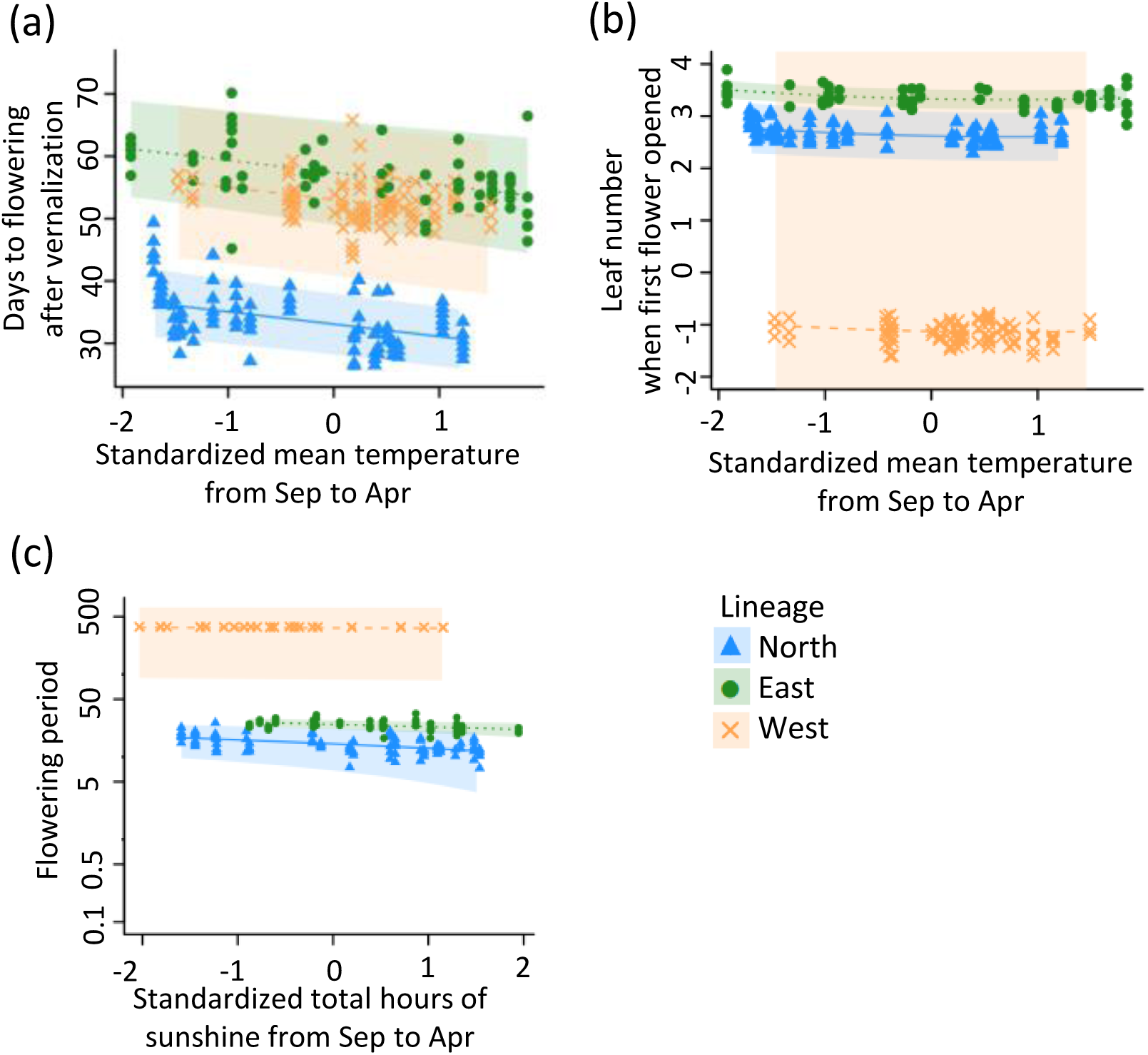
Partial dependence plots representing relationships which were common to all lineages. Each triangle, circle, and cross denotes an individual plant in the north, east, and west lineages, respectively. The solid, dotted, and dashed lines represent predicted relationships for the north, east, and west lineages and shaded areas represent confidence intervals. Predicted mean relationships and confidence intervals were calculated using the *lsmeans* package (Lenth 2006) in R by fixing the other parameters at their means. Note that because the values of the x axes are scaled to mean=0 and SD = 1, the slopes of the relationships represent their effect sizes.

## Discussion

Our common garden experiments demonstrated that the phenotypic variations in several traits of *C. hirsuta* have genetic bases determined partly by seed families and partly by lineages, genetic variation within each lineage, and climate factors. In particular, the effects of the lineages were strong for most of the traits. Moreover, a consistent effect of temperature on flowering phenology was detected; with increasing temperature in the seed source habitats, the timing of flowering became earlier in all three lineages. This result suggests that the genetic differentiations in response to temperature gradients have been constructed repeatedly in the non-native ranges encountered by *C. hirsuta*.

### Differentiation among the lineages and effects of multiple introductions

In our experiment, most of the phenotypic traits were significantly different among the lineages (Table 1). For example, the East lineage showed later flowering phenology and larger mass, whereas the West lineage showed earlier flowering phenology and smaller mass. One of the main causes of such phenotypic differences among the lineages could be differences among the introduced lineages. According to the results of ANOVA (Table 1), we could confirm that differences among lineages have caused large differences in some phenotypic traits. Evidence that multiple introductions provide genetic and phenotypic variation in introduced populations is accumulating (Bossdorf et al. 2005; Roman and Darling 2007; Xu et al. 2010; Tang et al. 2022), however, few studies have distinguished and quantified phenotypic variations among lineages of alien plants. In the case of *C. hirsuta* in Japan, we have succeeded in quantifying the phenotypic variations of distinct lineages due to this plant’s high rate of self-fertilization and early stage of invasion, and demonstrating that multiple introductions have increased the phenotypic variation in the non-native area. Unfortunately, there is no information about variation in traits in the source area that would enable us to evaluate the effects of multiple introductions more clearly.

### Phenotypic differentiation within each lineage and the similarity among the lineages

Among the relationships of genetic variation and climatic factors to the phenotypic variations we detected, three relationships were similar among the three lineages.

The effect of lower temperature on later flowering was detected most clearly among the three common relationships between plant traits and climate factors. The families from warmer areas tended to show earlier flowering and to have fewer leaves at the onset of flowering. Adaptation of flowering phenology to different temperature conditions has been repeatedly observed in spring-flowering species. Fitter and Fitter (2002) analyzed past and present data (in total four decades) of first flowering dates in 385 plant species including *C. hirsuta* and demonstrated that the first flowering dates of spring-flowering species became earlier with increased temperature due to climate change. In *Arabidopsis thaliana*, which is also a spring-flowering species and phylogenetically close to *C. hirsuta*, higher-altitude types had genetically later flowering phenology and larger mass than lower-altitude types (Montesinos-Navarro et al. 2011). Montesinos-Navarro *et al*. (2011) proposed a hypothesis that late flowering phenology is advantageous in environments with long, severe winters at high altitudes, whereas early phenology is advantageous in environments with short, mild winters at low altitudes. This is because, in environments with long, severe winters, plants are at the risk of damage by frost and sudden temperature decline in early spring. Later flowering is expected to help flowers and fruits to avoid such risk. By contrast, in environments with short, mild winters, plants are not likely to suffer from such risks, but rather to suffer stress from heat and drought in late spring. Rapid flowering after the end of winter is expected to help flowers and fruits to avoid such stress. This hypothesis seems to be also applicable to *C. hirsuta*. During range expansion, individuals with later flowering phenology may be able to survive in areas with severe winters and low temperatures. On the other hand, rapid flowering phenology may be advantageous in areas with mild winters and higher temperatures. Thus, for spring-flowering species, flowering phenology strategies aligned with the environment are critical for survival and reproduction.

As temperature of origin increased, not only days to flowering but also leaf number when the first flower opened decreased. At first glance, this result seems to be because early flowering may cause a decrease in the duration of leaf growth prior to flowering. A positive weak correlation was observed between the days to flowering and the leaf number (Pearson’s correlation coefficient = 0.26; *p* < 0.001) when all lineages were pooled for the analysis. However, when analyses within each lineage were conducted the correlation was not detected in the West lineage (Pearson’s correlation coefficient = -0.15; *p* = 0.15). This result indicates that replication using lineage helps to understand accurate correlations among traits and avoid misleading results from pooled data.

Strains from sites with more hours of sunshine tended to have shorter flowering periods. As *C. hirsuta* is an autogamous species, a long flowering period may not contribute to reproductive success. In our observations, insects that visited *C. hirsuta* during the flowering period were not pollinators but herbivores such as Aphidoidea. Although further field observations during flowering are needed, a shorter flowering period under sunny conditions might be effective for avoiding risks from herbivores and other stresses such as solar radiation, competition with another plants, or weeding.

The northwest area (facing the Japan Sea) of Japan is snow covered and has few hours of sunshine in winter. Although we detected a negative correlation between hours of sunshine and flowering period, there is a possibility that snow cover affected flowering period. Repressed growth under snow during winter might slow flower growth. Although the effects of snow on flowering phenology have been studied mainly in alpine plants (Kudo 1992; Wipf and Rixen 2010), widespread species also have the potential to be adapted to local snowfall.

We also found many relationships between climatic factors and plant traits that showed different patterns among the three lineages. One possible explanation for these observations is that differences in genetic and/or phenotypic variations within lineages lead to different responses to encountered environments. For example, the West lineage has lower genetic variation than the other lineages. If the lower variation was caused by bottleneck effects, it might have reduced the possibility of adaptation to new environments. Another possible explanation is the different climatic ranges encountered by the lineages; the nature and strength of the selective pressure may differ among the climatic ranges encountered. For example, cold may be a selective pressure on the North lineage, but it may have weak effects on survival in the West lineage. By clarifying the differences in responses among the lineages in reciprocal transplant experiments, we will be able to better understand the process of differentiation.

### Phenotypic differentiation during range expansion and Perspective

In this study, we narrowed down the candidates for responses that promote range expansion by exploring patterns common to the three lineages and showed both independent and parallel differentiations. Thus, we can observe replications of differentiation by distinguishing lineages having different origins. Such observations may be a novel utility of distinct lineages for the study of range expansion and evolution.

Our results suggested that rapid responses of plant traits are likely to result from natural selection in the new environmental conditions. To test this hypothesis, a reciprocal transplant experiment would be an effective approach. In such an experiment, we could examine whether the differentiations detected in this study actually contribute to increased fitness. In addition, as *C. hirsuta* has been studied well genetically (Gan et al. 2016; Hay and Tsiantis 2016), we would be able to utilize the accumulated knowledge of genetics to elucidate the genetic background of differentiation. These advanced studies could link the mechanisms in the ecological and genetic processes for the differentiation of traits, and provide a better understanding of range expansion of plant species.

## Acknowledgements

We thank Tomoyuki Itagaki, Yusuke Fusato and Masashi Sato for help with growth experiments, and Hideyuki Doi for his useful comments.

## Author Contributions

SM, HK and SS conceived and designed the experiments. SM, HK and SS conducted material sampling, and SM performed the experiments. SM and MO analyzed the data. All authors contributed to writing the manuscript.

